# Isolation and characterisation of quercitrin as a potent anti-sickle cell anaemia agent from *Alchornea cordifolia*

**DOI:** 10.1101/2021.11.11.468287

**Authors:** Olayemi Adeniyi, Rafael Baptista, Sumana Bhowmick, Alan Cookson, Robert Nash, Ana Winters, Luis A. J. Mur

## Abstract

*Alchornea cordifolia* Müll. Arg. (commonly known as Christmas Bush) has been used traditionally in Africa to treat sickle cell anaemia (a recessive disease, arising from the *S* haemoglobin [Hb] allele) but the active compounds are yet to be characterised. Herein we describe the use of sequential fractionation coupled with *in vitro* anti-sickling assays to purify the active component. Sickling was induced in Hb*SS* genotype blood samples using sodium metabisulphite (Na_2_S_2_O_5_) or incubation in 100 % N_2_. Methanol extracts of *A. cordifolia* leaves and its sub-fractions showed >70 % suppression of HbSS erythrocyte sickling. Purified compound demonstrated 87.2 ± 2.39 % significant anti-sickling activity and 93.1 ± 2.69 % erythrocyte sickling-inhibition at 0.4 mg/mL. Nuclear magnetic resonance (NMR) spectra and high-resolution mass spectroscopy identified it as quercitrin (quercetin 3-rhamnoside). Purified quercitrin also inhibited the polymerisation of isolated HbS and stabilized sickle erythrocytes membranes. Metabolomic comparisons of blood samples using flow-infusion electrospray-high resolution mass spectrometry indicated that quercitrin could convert HbSS erythrocyte metabolomes to be similar to HbAA. Sickling was associated with changes in anti-oxidants, anaerobic bioenergy and arachidonic acid metabolism, all of which were reversed by quercitrin. The findings described could inform efforts directed to the development of an anti-sickling drug or quality control assessments of *A. cordifolia* preparations.

## Introduction

Sickle cell anaemia (SCA) is an autosomal recessive genetic blood disorder arising from the *S* allele of haemoglobin (Hb). SCA is particular prevalent in the tropics and especially in sub-Saharan Africa where the *S* allele may confer to some, tolerance to malaria. The World Health Organization (WHO) estimates that 300,000 children are born with SCA annually, 75% of whom are in sub-Saharan Africa (Roucher *et al*., 2012).

SCA arises from a single amino acid substitution of glutamic acid with hydrophobic valine in the Hbβ-globin chain. This results in an altered haemoglobin tetramer (α2βs2), haemoglobin S (Hb S) (Igwe *et al*., 2017; Olubiyi *et al*., 2019; Sundd *et al*., 2019). HbS polymerises under hypoxic conditions, due to hydrophobic interactions between β6Val on two deoxy-HbS molecules (Rees *et al*., 2010). The polymer of helical fibres lengthen and stiffen to cause the characteristic sickle shape of HbSS erythrocytes (Higgs, 1986; Edelstein *et al*., 1973; Odievre *et al*., 2011). Polymerization is linked to a dysregulation of cation homeostasis resulting from the activation of some ion channels, particularly the K^+^/Cl^-^ co-transport system and the Ca^2+^-dependent K^+^-channel (Gardos channel). Ca^2+^ activation of the Gardos channel, increases H_2_O and K^+^ efflux leading to the dehydration of sickled erythrocytes (Brugnara, 1995). Haemoglobin is denatured to form hemichromes; histidine-linked complexes, on the internal surface of the membrane. The haem group releases Fe^3+^ to foster an oxidizing microenvironment (Odièvre et al., 2011).

The effects of SCA can be mitigated through episodic blood transfusions to stabilize the Hb levels (Friedman *et al*., 2018; Piccin *et al*., 2019), increasing the provision of oxygen (Piccin et al., 2019), and rehydration with intravenous fluids (Carden *et al*., 2019; Peterfy *et al*., 2018). The pain associated with SCA crises can be managed with nonsteroidal anti-inflammatory drugs (NSAIDs) or other non-opioid analgesics (Lakkakula *et al*., 2018; Puri *et al*., 2018). However, there are relatively few chemical agents which interfere with the mechanism and/or kinetics of the sickling process, hydroxyurea and voxelotor being most often used (Abdulmalik *et al*., 2005; Omar *et al*., 2019). Such therapies have attendant limitations (Best *et al*., 1998; FDA, 2019; Rees *et al*., 2010) such as high costs and low availability for millions of patients in sub-Saharan Africa (Kato *et al*., 2018) as well as attendant risk factors with long term clinical use (Friedman *et al*., 2018; Imaga, 2010; Odunlade *et al*., 2017; Oyewole *et al*., 2008). There is a need for new cost-effective anti-sickling small molecules to treat SCA.

Medicinal plants are widely used in Sub-Saharan Africa to manage SCA and have driven research into their active components. Thus, phenylalanine and p-hydroxy benzoic acid (PHBA) from *Cajanus cajan* (.Akojie & Fung, 1992), zanthoxylol (Sofowora et al., 1975), betulinic acid (Tshibangu, 2010), divanilloylquinic acids from *Fagara zanthoxyloides* Lam. (Rutaceae) (Ouattara *et al*., 2009), butyl stearate from *Ocimum basilicum* (Tshilanda *et al*., 2014) and ursolic acid from *Ocimum gratissimumL*. (*Lamiaceae*) (Tshilanda et al., 2015) have been linked to reduced sickling. Leaves of *Alchornea cordifolia* are used as a ‘blood tonic’ to reduce the symptoms of SCA in Nigeria. It has featured in some research that have characterised its biochemistry (Borokini & Omotayo, 2012; Ogundipe, 1999; Oloyede *et al*., 2010; Schmelzer, 2007; Mohammed *et al*., 2013; Ogundipe *et al*., 2001) but the active compounds linked to anti-sickling activities have not been defined.

In this paper, we describe the isolation and characterisation of quercitrin from the Nigerian shrub, *A. cordifolia* as the main anti-sickling agent. Quercitrin was able to prevent and reverse *in vitro* HbSS erythrocyte sickling, primarily through inhibition of HbS polymerization and membrane stabilization under hypoxic conditions.

## Materials and Methods

### Chemicals

LC-MS-grade formic acid in water and acetonitrile and HPLC-grade solvents; methanol, dichloromethane, acetonitrile, ethyl acetate, and *n-*hexane and Nitrogen (N_2_) were all obtained from Fisher Scientific, Leicestershire, (UK). Sodium metabisulfite (Na_2_S_2_O_5_), rutin, gallic acid, linoleic acid, 1,1-diphenyl-2-picrylhydrazyl (DPPH), ascorbic acid, quercetin, 2,2′-azobis(2-amidinopropane) dihydrochloride (AAPH), phosphate-buffered saline (PBS) pH 7.4, *p*-hydroxybenzoic acid (PHBA), L-phenylalanine, sodium chloride (NaCl), trifluoroacetic acid (TFA), Triton X-100 sodium dihydrogen phosphate, sodium chloride and L-phenylalanine were all purchased from Sigma-Aldrich (UK).

### Collection of plant samples

*A. cordifolia* leaves were harvested from around Ado-Ekiti, South-West, Nigeria in May, 2015. Samples were deposited as a voucher specimen (UHAE2020030) at a herbarium, Department of Botany, Ekiti State University, Nigeria. The leaves were shade-dried for 3 weeks until completely dehydrated.

### Blood sampling

A HbSS blood sample from a single clinically diagnosed SCA sufferer (the first author) was used to evaluate the anti-sickling activities of the plant extracts. 5 mL of blood were extracted using a lavender topped vacutainer (BD Vacutainer tubes, UK) which uses dipotassium/tripotassium salts of EDTA as an anticoagulant. A control HbAA sample was taken from a single volunteer (the corresponding author). The whole blood (4 mL) was centrifuged at 2800 x g for 10 min at 4 °C to sediment the erythrocytes. The plasma supernatant was removed, and erythrocytes underwent sequential centrifugations (2800 g for 10 min at 4 °C) involving washing three times in phosphate buffered saline (PBS) pH 7.4. Cells were finally resuspended in 5 volumes PBS to 1 packed volume of erythrocytes and used immediately.

### Extraction of *Alchornea cordifolia* leaves

Pulverized air-dried leaves of *A. cordifolia* (1 kg) was extracted 12.5 L dichloromethane (CH_2_Cl_2_; DCM) and then sequentially with 51.5 L 75 % methanol (75 % MeOH; ALM) and two rounds of 10 L of sterile deionised water (H_2_O; “Aqueous”, “Aqueous2”) at room temperature with constant stirring. Each extraction occurred for a 72 h period. After filtration, the extracts were concentrated under reduced pressure at 40 °C and stored at −20 °C until further use. The aqueous extract was freeze-dried.

### Bioactivity-guided purification of MeOH extract

Partial purification of MeOH extract (ALM; 96.5 g) involved using a modified method as described by Rani & Devanand, 2011 (Fig. S1). The crude extract of *A. cordifolia* was separated using silica gel (243.43 g as adsorbent, 70–200 mesh, Material Harvest, UK) packed in a 40 × 500 mm chromatographic column and eluted with a continuous solvent gradient of increasing polarity; ethyl acetate (EtOAc) (0–100 %) in hexane and then methanol (0–100 %) in EtOAc. According to differences in composition (as indicated by thin layer chromatography), 19 fractions were obtained (ALM1–ALM19)). Thin layer chromatography (TLC) was performed on Sigma-Aldrich silica gel 60 F_254_ gel plates and visualized under UV light and by spraying sulphuric acid-MeOH (1:1) followed by heating.

Fractions 7 (ALM7) and 8 (ALM8) were combined and separated to yield 23 sub-fractions (ALM7A-ALM7W) on a silica gel-packed chromatographic column (70–200 mesh) and eluted with a solvent gradient of increasing polarity; ethyl acetate (0–100 %) in hexane and then methanol (0–100 %) in ethyl acetate.

Following anti-sickling assays ALM7T was further separated by preparative HPLC using a C18 3.5 μm, 4.6 × 50 mm column (Waters, UK). The eluting gradient was as follows 90 % water 10 % acetonitrile for 2.5 min then going to 100 % acetonitrile at 8.5 min and continuing at 100 % until 13min. There was 0.01 % trifluoroacetic acid present throughout and the flow rate is 1.5 mL/min. This yielded 8 peaks separated into single fractions, ALM7T1-ALM7T8.

### Ultra high performance liquid chromatography–high resolution mass spectrometry (UHPLC-HRMS)

Fractions were analyzed on an Exactive Orbitrap (Thermo Fisher Scientific) mass spectrometer, which was coupled to an Accela Ultra High Performance Liquid Chromatography (UHPLC) system (Thermo Fisher Scientific). Chromatographic separation was performed on a reverse phase (RP) Hypersil Gold C18 1.9 μm, 2.1 × 150 mm column (Thermo Scientific) using H_2_O using 0.1 % formic acid (v/v, pH 2.74) as the mobile phase solvent A and ACN: isopropanol (10:90) with 10 mM ammonium acetate as mobile phase solvent B. Each sample (20 μL) was analysed using 0-20 % gradient of B from 0.5 to 1.5 min and then to 100% in 10.5 min. After 3 min isocratic at 100 % B the column was re-equilibrating with 100% A for 7 min.

### Nuclear Magnetic Resonance

NMR spectra were obtained using Bruker Ultra shield-500 NMR spectrophotometer (1H-NMR 500 MHz, 13C-NMR 100 MHz) using either MeOD as the solvent reference.

Quercitrin (2-(3,4-dihydroxyphenyl)-5,7-dihydroxy-3-[(2*S*,3*R*,4*R*,5*R*,6*S*)-3,4,5-trihydroxy-6-methyloxan-2-yl]oxychromen-4-one) (Fig. 1): Yellow powder; *m/z* 447.09363 [M+H]^+^ (calcd. For C_21_H_20_O_11_, 448.100561). ^1^H-NMR (500MHz, MeOD): *δ* 0.95 (3H, d, *J*=6.0 Hz), 3.32 (1H, m), 3.43 (1H, m), 3.76 (1, dd, J=3.0 and 3.0 Hz), 4.23 (1H, s), 5.36 (1H, s), 6.20 (1H, d, *J*=1.8 Hz), 6.37 (1H, d, J=1.8 Hz), 6.91 (1H, d, J=8.4 Hz), 7.30 (1H, dd, J=8.4 and 2.1 Hz), 7.34 (1H, d, J=2.0 Hz) ppm. ^13^C-NMR (100 MHz, MeOD): *δ* 17.67, 41.90, 71.96, 72.05, 72.24, 73.38, 94.79, 99.88, 103.58, 106.00, 116.45, 117.07, 122.95, 123.10, 136.31, 146.42, 149.79, 158.56, 159.35, 163.22, 165.83, 179.70 ppm. These spectroscopic data were in agreement with previous studies for quercitrin (Jang *et al*., 2011; Zhong *et al*., 1997).

**Fig. 1.**
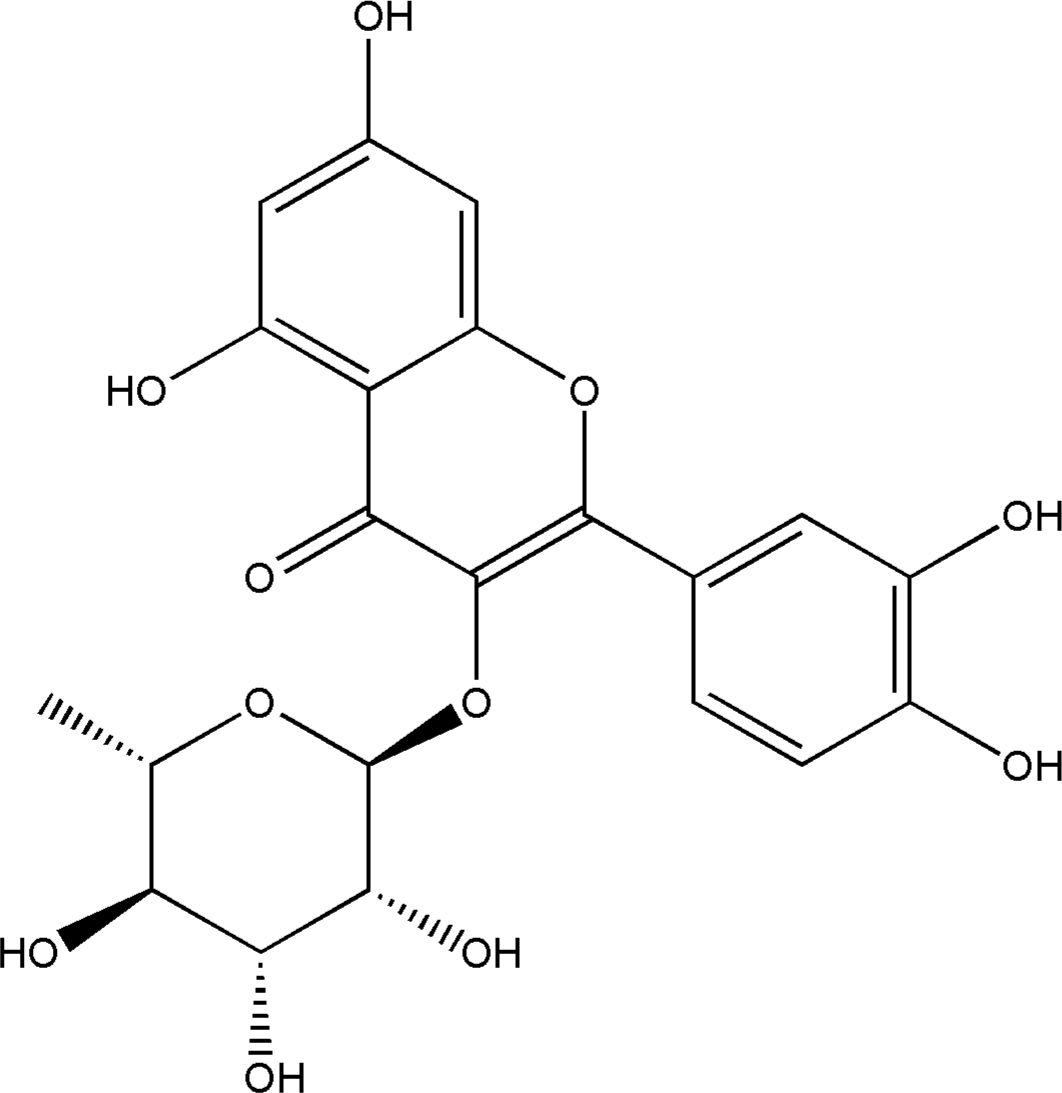
The chemical structure of Quercitrin

### *In vitro* sickling-inhibition and reversibility assays

The sickling-inhibition assay consisted of 100 μL of HbSS erythrocytes, 100 μL of PBS, and 100 μL of the test extract and incubated at 37 °C for 2 h. To induce sickling, freshly prepared 2 % (w/v) Na_2_S_2_O_5_ solution (300 μL) were incubated with the cells for an additional hour in a water bath at 37 ºC. Cells were then fixed with 3 mL of 5 % (w/v) buffered formalin solution. A total of 10 μL of the incubated cells were transferred to a haemocytometer and five fields counted on each slide using a Leica ATC 2000 Binocular Phase Contrast Microscope at 40 x magnification. Cells were classified as either normal or sickled. Each assay was repeated five times to generate the data presented.

For reversibility assays, cells were prepared as above and incubated with 2 % (w/v) Na_2_S_2_O_5_ at 37 °C for 1 hour. Then, 100 μL of each sample was added and incubated at 37 °C for an additional 4 h period. Cells were fixed and mounted on a haemocytometer and counted as described earlier.

### Erythrocyte leakage assay

Sterile microcentrifuge tubes were each filled with 1 mL HbSS erythrocytes and centrifuged at 2,800 x g for 10 min at 4 °C. The supernatant was discarded, and the erythrocytes were washed three times with phosphate buffered saline (PBS; 0.01 M pH 7.4) and resuspended in 4 % v/v PBS. 90 μL was added to wells on a 96 well plate and 10 μL of each sample being tested (at 10 x final concentration) was added to wells of the first row and then serial-dilutions of the sample down to row 6. Row 7 contained erythrocytes suspensions with 0.1 % Triton-X100 (Sigma-Aldrich UK) as a positive control and row 8 contained only erythrocytes as the negative control. The plate was incubated for 1 h at 37 °C. Following a 5 min centrifugation at 2,800 x g at 4 °C, 70 μL of supernatant from each well was transferred to a transparent, flat bottom 96 well plate. Changes in absorbance (OD_450_nm), indicating haemoglobin leakage were measured using the Hidex Sense Plate Reader (LabLogic, Sheffield UK). This absorbance was used to calculate percentage haemolysis (0.1 % triton X-100 = 100 % and the negative control = 0 %). Experiments were performed in quadruplicate.

### Hb polymerisation assay

The method of (Iwu et al., 1988) was adapted for 96-well plates to assess polymerized Hb SS turbidity. A haemolysate was prepared by adding 2 mL of ice-cold distilled water to packed, washed erythrocytes and then the cellular debris was pelleted by centrifugation at 6,000 x g for 20 min at 4 °C. Assays of 220 μL 2 % Na_2_S_2_0_5_, 20 μL of the test compound (at 5 different concentrations and using PBS as a control) and 50 μL HbS haemolysate (1:5 v/v dilution in PBS) was incubated in a 96-well plate. This was shaken and the absorbance at 700 nm taken in 30 second intervals for period 20 minutes (Hidex sense microplate reader, LabLogic, Sheffield, UK). Tests were carried out in quadruplicate.

### Scanning Electron Microscopy (SEM)

Erythrocytes were fixed in 2 % glutaraldehyde PBS for 30 min, and rinsed three times in 0.075 M sodium phosphate buffer (pH 7.4). Samples were then incubated with 2 % OsO_4_ in PBS (pH 7.4) for 2 h at 4 °C and then rinsed thrice in 0.075 M sodium phosphate buffer (pH 7.4). Subsequently, the sample was dehydrated in 30 %, 50 %, 70 %, 90 %, and finally, three changes of 100 % ethanol. For SEM, 200 μL of the samples in hexamethyldisilazane (HMDS) were air dried on coverslips, coated with carbon and imaged using a Zeiss Ultra plus FEG SEM.

### Metabolomic analyses

Samples of 500 μL of washed HbSS erythrocytes, 500 μL of PBS and 500 μL of quercitrin (0.5 mg/ml in final volume) were mixed and incubated at 37 °C for 2 h. Sickling was induced through chronic deoxygenation; the reaction mixture (1500 μL) in an anaerobic tube was deoxygenated by gently bubbling 100 % N_2_ through in a 37 °C water bath for 12 h. After the incubation period, the morphology of the cells was confirmed using light microscopy.

Erythrocyte extractions were carried out using published protocols (Sana *et al*., 2008; Srivastava *et al*., 2017). Extracts were transferred to a 2 mL microcentrifuge tube and this was dried using a SpeedVac at 4 °C. The pellets were then resuspended in 100 μL 50 % methanol, in a HPLC vial containing a 0.2 mL flat bottom micro insert for flow infusion electrospray ion high resolution mass spectrometry (FIE-HRMS) analysis. FIE-HRMS was performed with an Exactive HCD mass analyser equipped with an Accela UHPLC system (Thermo-Scientific). Data acquisition for each individual sample was conducted in alternating positive and negative ionisation mode, over four scan ranges (15-110, 100-220, 210-510, 500-1200 *m/z*) with an acquisition time of five min. Individual metabolite *m/z* values were normalised as a percentage of the total ion count for each sample. Data were normalised to total ion count and log_10_-transformed. Metabolites and pathway identification were performed by the MetaboAnalyst 4.0. MS peaks to pathway algorithm (Chong et al., 2018) (tolerance = 5 ppm, reference library; *Homo sapiens*). This involved metabolites being annotated using KEGG database, considering the following possible adducts: [M]^+^, [M+H]^+^, [M+NH_4_]^+^, [M+Na]^+^, [M+K]^+^, [M-NH_3_+H]^+^, [M-CO_2_+H]^+^, [M-H_2_O+H]^+^; [M]^−^, [M−H]^−^, [M+Na−2H]^−^, [M+Cl]^−^, [M+K−2H]^−^. For each mass-ion (*m/z*) the annotation was made using a 5 ppm tolerance on their accurate mass and the different adducts formed for each metabolite.

### Statistical analysis

The statistical analyses used SPSS version 26.0 software and XLSTAT_2020.1.1.64525. One Way Analysis of Variance (ANOVA) coupled with the Tukey’s *post-hoc test* was used to compare the data and to identify means with significant differences. *P* values of <0.05 were considered significant.

## Results

### Sample extraction and anti-sickling activity of *A. cordifolia* crude extracts

Sequential extractions using DCM, 75 % MeOH, and 100 % water from the dried leaves, derived samples of 2.47, 9.65, and 8.28 % dry weight. The sickling-inhibitory activities of *A. cordifolia* leaf-extracts were compared at different concentrations (1, 5 and 10 mg/mL in PBS). The proportion of sickled erythrocytes could be readily assessed on a haemocytometer (Fig. S2 A, B and C) and Fig. 2A, B show the data obtained for 1 mg/mL of extract. The MeOH extract exhibited significantly (*P*<0.01) higher sickling-inhibitory activity (91.4 %) than any other extract (Fig. 2A, B). MeOH extract also exhibited a better sickling reversibility activity than any other extract (Fig. 2C).

**Fig. 2.**
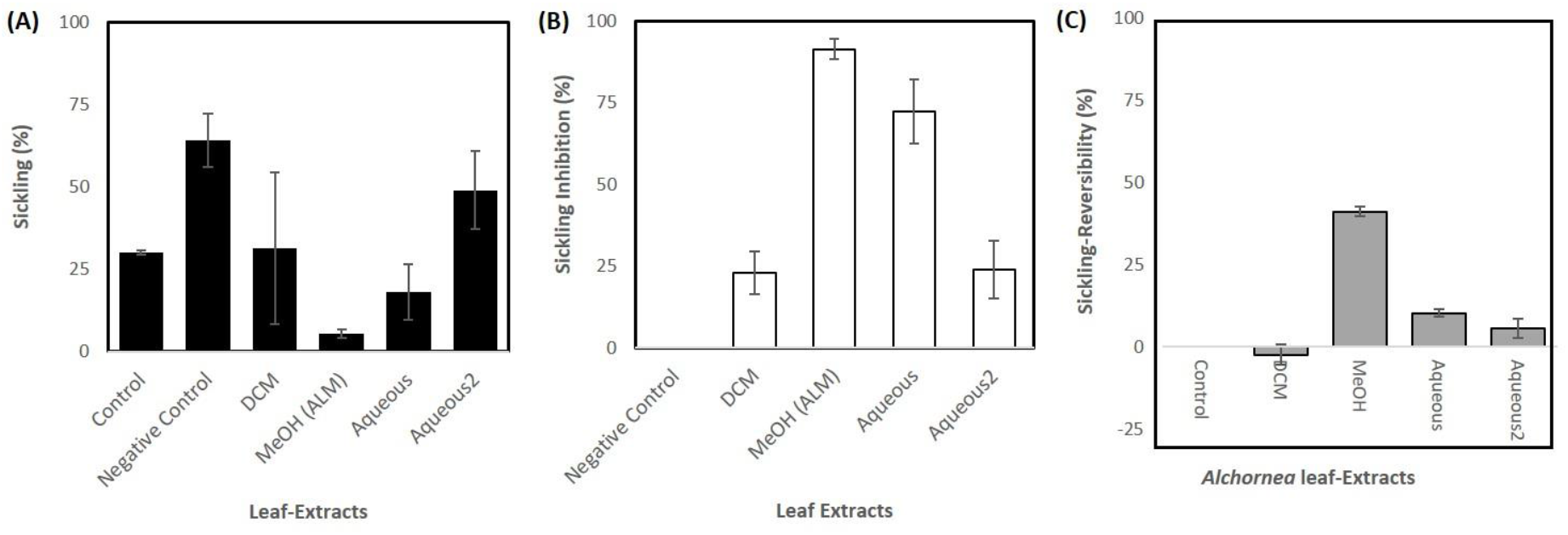
The effect of *Alchornea spp* leaf-extracts (1 mg/mL) on percentage sickling on incubation with erythrocytes in Na2S2O5-induced hypoxic condition. (A) Percentage sickling (B) Percentage sickling-inhibition (C) percentage reversion of sickling. Data represent the average of 3 similar results from repeat experiments.

The cytotoxicity of aqueous and MeOH leaf-extracts of *A. cordifolia* on HbSS erythrocytes was evaluated using an erythrocyte leakage assay (Fig. S3). Some haemolysis was observed in both MeOH and aqueous extracts at 10 mg/mL but this was only <2 % at 1 mg/mL.

### Isolation of the anti-sickling bioactives in the *A. cordifolia* methanolic extract

A bioassay-guided purification process was used to isolate the active bioactives in the MeOH extract of *A. cordifolia* (ALM; 50 g) (Fig. S4). Of a total of 19 fractions, three fractions ALM5 (0.2 % yield), ALM7 (0.6 % yield) and ALM8 (5.3 % yield) exhibited sickling inhibition above 95 % at 1 mg/mL concentration; ALM7 (96.5 ± 2 .8 %) and ALM8 (96.7 ± 2.7 %). ALM7 and ALM8 were combined due to their chemical similarities as revealed by TLC. A total of 23 sub-fractions (ALM7A-ALM7W) were fractionated from combined ALM7 and ALM8. As enrichment of the bioactive product was expected following fractionation, sickling-inhibiting activities were assessed at a lower concentration range than previously (Fig. S5). Sickling-inhibiting activities of >70 % were observed with 0.25 mg/mL in sub-fractions ALM7N, ALM7Q, ALM7T and ALM7V.

Sub-fraction ALM7T was further separated by preparative-HPLC into individual compounds to yield ALM7T1-ALM7T8. Screening for anti-HbSS erythrocyte sickling properties indicated that ALM7T5 showed > 70 % sickling-inhibition activity at 0.5 and 0.25 mg/mL (Fig. 3). Representative haemocytometer images can be seen in Fig. S2. NMR was used to identify the only chemical detected within these peaks. ALM7T5 was identified as quercitrin (quercetin-3-rhamnoside), a flavonol glycoside, based on high resolution LC-MS and NMR data.

**Fig. 3.**
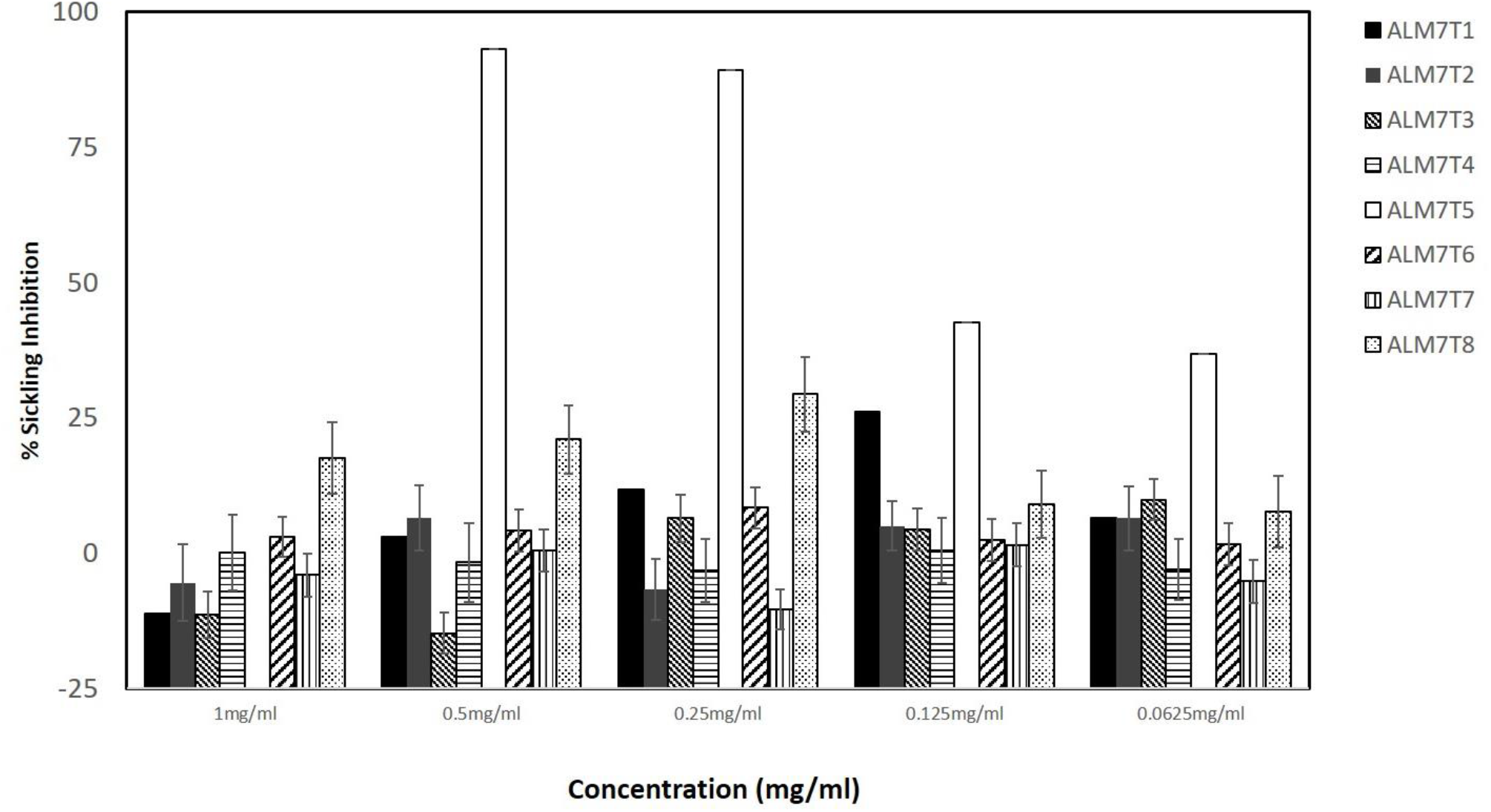
The effect of fraction ALM7V on *ex vivo* erythrocyte-sickling in Na_2_S_2_O_5_-induced hypoxia.

The ability of quercitrin to reverse sickling in erythrocytes in Na_2_S_2_O_5_-induced hypoxia after a 4-h incubation period was tested. The highest concentration tested, 0.80 mg/mL, could reverse sickling (41.8 % ± 4.8) and activity was still detected (18.1 ± 1.2) at 0.05 mg/mL (Fig. 4).

**Fig. 4.**
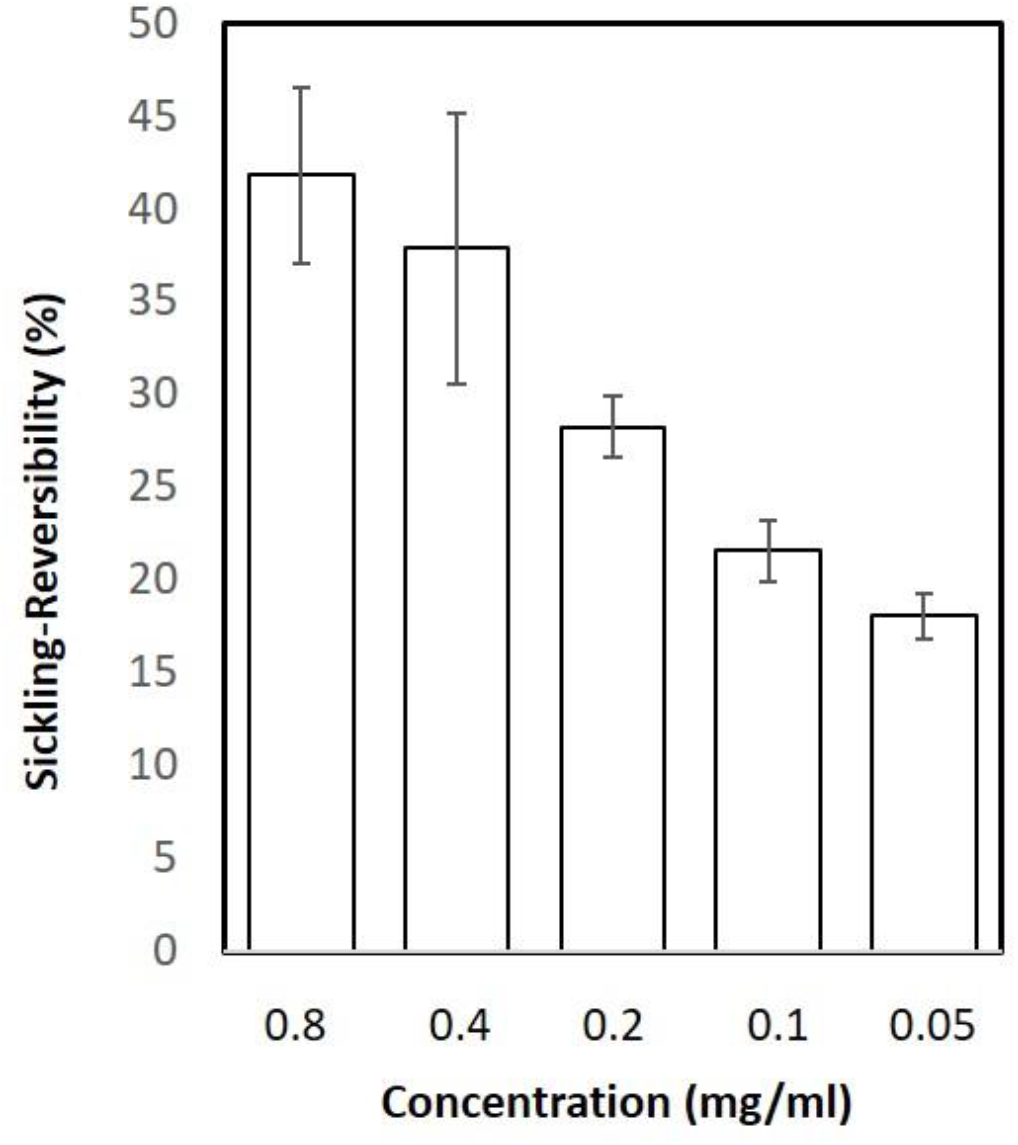
Reversibility effects of Quercitrin on HbSS-RBC sickling *in vitro*, at low oxygen tension induced by Na_2_S_2_O_5_ after a total 5 hour-incubation period. Data represented as mean and SD, obtained from three typical independent experiments.

The biological activity of quercitrin has been previously associated with its aglycone, quercetin (Shen *et al*., 2003; Xiao, 2017). Therefore, the anti-sickling properties of quercetin and quercitrin were compared across a concentration range (Fig. S6). The results showed significantly (*p* < 0.01) higher anti-sickling activity (ranging between 93.1 % ± 1.6 to 36.9 % ± 1.9 %) in quercitrin compared to quercetin (ranging between 11.8 % ± 0.98 to -1.14 % ± 1.0). This suggested that quercitrin, but not quercetin, was able to decrease erythrocyte sickling.

### Quercitrin exhibits anti-HbSS polymerisation activity

Different concentrations of quercitrin (0.25-4 mg/mL) were tested for its ability to inhibit the polymerisation of deoxygenated HbS over 20 min (Fig. 5). Deoxy HbSS and HbAA haemolysates (with Na_2_S_2_O_5_- and phosphate buffer-incubation) were used as controls. Quercitrin prevented HbS polymerisation over all tested concentrations so that there was no significant difference to HbAA results (Fig. 5A). Comparison was also made with PHBA, known to inhibit sickling in erythrocytes (Atabo *et al*., 2016) (Fig. 5B). PHBA showed similar ability to suppress HbSS polymerisation exhibited but at 0.5 and 0.25 mg/mL, suppression was significantly (*p >* 0.05) less effective than quercitrin (Fig. 5B). Quercetin exhibited no ability to prevent the polymerisation of HbSS with no significant difference to the positive control (Fig. S7). Taken together, these data indicate that quercitrin is a potent inhibitor of *in vitro* HbS polymerisation and this is most likely to represent its main mode of action in inhibiting and reversing HbSS-erythrocyte sickling.

**Fig. 5.**
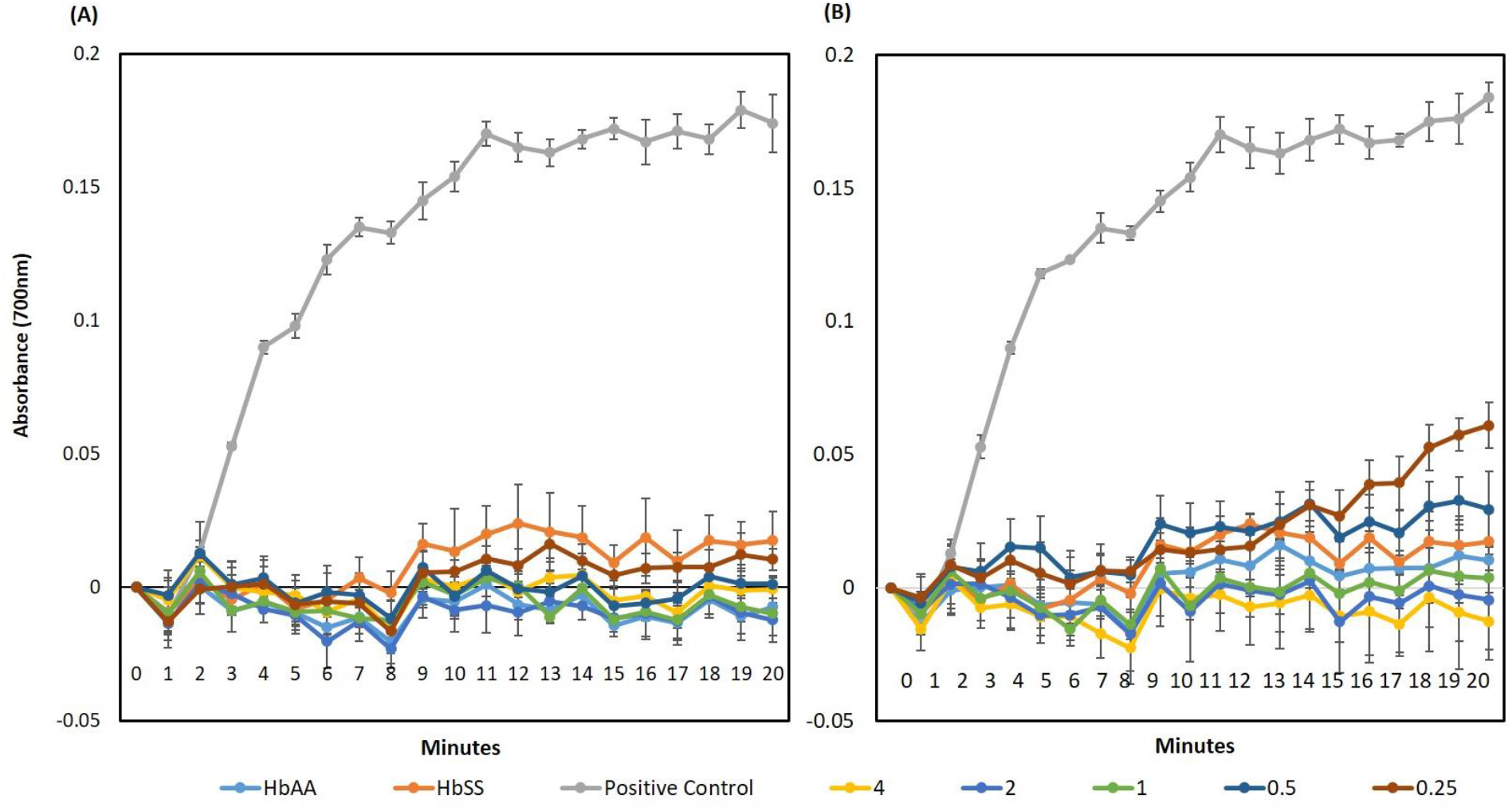
Inhibitory Effects of different concentrations of 0.25-4 mg/mL of **(A)** quercitrin and **(B)** p-hydroxybenzoic acid on HbS polymerisation, *in vitro*. Quercitrin inhibits HbS polymerisation compared to the “positive control”: deoxyHbS without quercitrin. With HbSS; HbS not subjected to deoxygenation. DeoxyHbA (HbAA) did not polymerise and represents a negative control. This represents data obtained from three typical independent experiments performed in quadruples.

### Assessing the impact on quercitrin on HbSS sickling using metabolomics

In using metabolomics to assess quercitrin’s mode of action, it was not possible to use 2 % Na_2_S_2_O_3_ to induce the sickling phenotype, as this would dominate the subsequent metabolite profile. Thus, an alternative approach was employed where anoxic conditions were induced through replacement of air with 100 % N_2_. This proven to be a highly successful approach as demonstrated using SEM (Fig. S8). Imposition of N_2_–induced anoxia led to sickling in HbSS erythrocytes but not HbAA erythrocytes (data not shown). HbSS erythrocytes treated with 0.5 mg/mL quercitrin exhibited morphologies which were similar but not identical to HbAA erythrocytes. Interestingly, erythrocytes treated with 0.5 mg/mL PHBA did all exhibit the normal discoid phenotype with some sickled cells observed. In the metabolomic treatments,

HbSS erythrocytes were either 1) maintained under normoxic conditions (SS), 2) deoxygenated with N_2_ (SS-N) only or 3) incubated with quercitrin followed by deoxygenation with N_2_ (SS-Q). Controls consisted of HbAA erythrocytes (AA) with no quercitrin treatment (Fig. 6). PCA indicated that the different experimental classes formed two clusters; one broadly assocated with sickled cells the other with non-sickled cells (Fig. 6A). SS and SS-N samples were both closely clustered across principal component 1 (PC1) which describes the major source of variation. This suggested pre-existing metabolomic changes in the HbSS erythocytes, even under normoxia. However, by adding quercitrin the hypoxic HbSS metabolome shifted change so that samples clustered with the HbAA group.

**Fig. 6.**
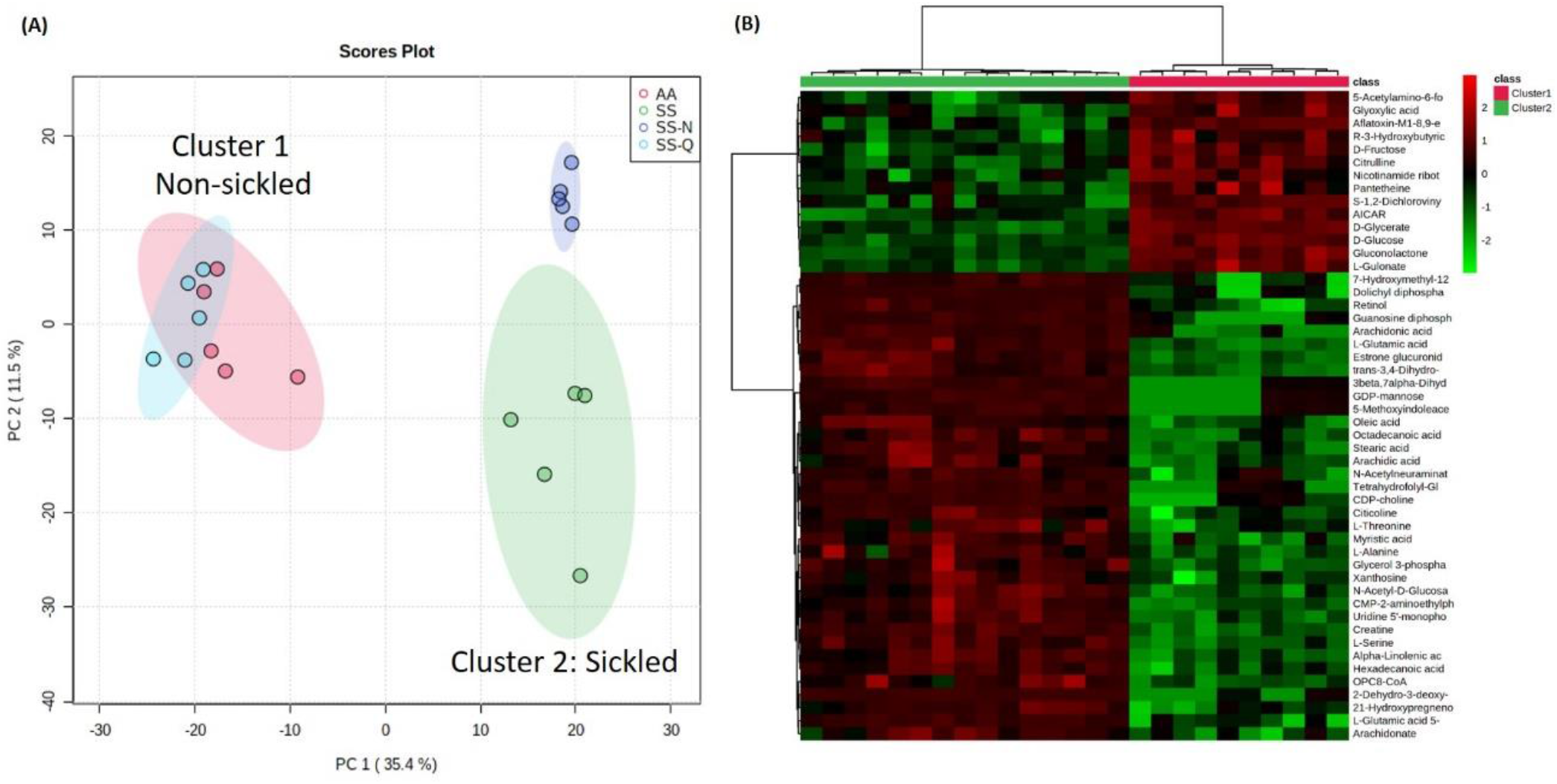
Principal Component Analysis and Hierarchical cluster Analysis the metabolomic impact of quercitrin on erythrocyte sickling. HbAA (AA) and HbSS (SS) erythrocytes were maintained under normoxic conditions or under N_2_ (N) and in some cases treated with quercitrin (0.5 mg/mL) (SS-Q) imposition of hypoxic conditions. **(A)** PCA of derived metabolite profiles for each treatment (note the separation of the samples into two main clusters reflecting sickled and non-sickled groups) **(B)** heatmap based on significant metabolite differences between the non-sickled (cluster 1) and sickled (cluster 2) groups

The sources of variation between sickled and non-sickled erythrocyte samples were determined (Fig. 5B). We could annotate 126 metabolites using KEGG database and their relative abundances discriminated between the sickled and non-sickled phenotypes as shown using a heatmap. The sickled group showed relative increases in metabolites such as arachidonic, stearic, myristic and linolenic acids which were suggestive of lipid processing. Sickled cells also appeared to be relatively deficient in glucose and fructose. These effects were reversed by quercitrin treatment. To provide functional information for these differences, biochemical enrichment analyses were conducted. The over-representation analysis (ORA) method to evaluate pathway-level enrichment based on significant features (*p*-value is measured with Fisher’s exact test) (Li *et al*., 2013), combined with gene set enrichment analysis (GSEA) which extracts biological meaning from a ranked metabolite list (Subramanian et al., 2005). A total of 6 metabolic pathways were identified to be significantly enriched (Table S1). These suggested that the metabolomic switches in the erythrocyte between the sickled and non-sickled stated apparently involves redox changes (ascorbate), thiol metabolism (cysteine), fatty acid processing and haem biosynthesis (porphyrin).

## Discussion

The high prevalence of SCA in the developing nations of West Africa is driving the requirement for cost-effective means to treat the disease. In Sub-Saharan Africa, medicinal plants are used widely to manage SCA, although, relatively few have been validated scientifically. In the case of *A. cordifolia*, an aqueous extract showed anti-sickling activity (Mpiana, Mudogo, *et al*., 2007; Mpiana, Tshibangu, *et al*., 2007) but the bioactives were not previously defined. In this study, we followed a bioactivity guided fractionation and purification strategy to define the quercitrin as the main active anti-sickling agent in *A. cordifolia*.

### Inhibition of HbS polymerisation: An important mode of action for quercitrin

Beyond simply defining quercitrin as the anti-sickling chemical, we considered its mode of action. Quercitrin has been previously demonstrated to have a low toxicity profile on mouse macrophages (IC_50_ of approximately 0.1mg/mL) (Muzitano et al., 2006) and have potent antioxidant (Bose *et al*., 2013; Yamazaki *et al*., 2007; Yin *et al*., 2013), antiapoptotic, (Sánchez de Medina *et al*., 1996; Yin *et al*., 2013), anti-leishmanial (da Silva *et al*., 2012; Wagner *et al*., 2006) anti-diarrhoeal (Galvez et al., 1993), anti-nociceptive (Gadotti et al., 2005) and anti-inflammatory activities (Camuesco *et al*., 2004; Gadotti *et al*., 2005; Muzitano *et al*., 2006). The latter include intestinal inflammatory diseases in experimental models of rat colitis (Camuesco *et al*., 2004; Cruz *et al*., 1998) and colonic inflammation (Cruz *et al*., 1998; de Medina *et al*., 1996; De Medina *et al*., 2002; Galvez *et al*., 1997). These anti-inflammatory effects may be mediated through the inhibition of the NF-κB pathway (Comalada et al., 2005).

Many of anti-sickling drug leads effect the Hb gene (Pule et al., 2015), the HbS molecule or are erythrocyte membrane modifiers (Mehanna, 2001). Our methods did not assess Hb gene modification, but we provide clear evidence that quercitrin affected the HbS protein to suppress polymerisation. This was important as clinically, the delay of HbS polymerisation during transit of erythrocytes through post-capillary venules is necessary for SCA disease modification (Ilesanmi, 2010). Our assessments of extracted Hb polymerisation, showed the expected results for the controls; i.e. no polymerisation with HbA (HbAA), oxygenated HbS (HbSS) but deoxyHbS did polymerise. However, with quercitrin, deoxyHb polymerisation was suppressed in a concentration-dependent manner comparable to PHBA. Crucially, a recent study used multi-spectroscopic and molecular simulation techniques show that isoquercitrin had anti-sickling activity through a direct interaction with haemoglobin molecules (Syed *et al*., 2019). Although not assessed in this paper, it is very probable that quercitrin has this property. Thus, quercitrin could act by stereospecific covalent or non-covalent attachment to the HbS molecule as an in HbS (Oder et al., 2016). Additionally, quercitrin was able to partially reverse erythrocytes sickling after a two-hour incubation period, suggesting quercitrin was able to modify already polymerized HbS and thus, have some ability to reverse polymerisation. This partial effect may reflect the rate at which quercitrin is transported across the membrane (Imaga, 2013).

The pharmacological activities of a given bioactive compound are dependent on its chemical structure (Heim et al., 2012). Previous investigations have credited the biological activity of quercitrin to the aglycone, quercetin (Shen et al., 2003; Xiao, 2017) but the possibility of the glycosidic residue was being crucial for its effects as glycoside activity has also been suggested (Křen, 2008; Xiao, 2017). The quercetin aglycone or glucoside is not found in human plasma, however conjugates, such as quercetin-3-glucuronide, quercetin-3’-sulfate, and isorhamnetin-3-glucuronide have been found (Janisch et al., 2004). It is generally thought that the flavonoid glycosides such as quercitrin, enters the colon and are hydrolysed to the aglycone by quercitrinase found in Enterobacteria (Tranchimand et al., 2010). The aglycone is then absorbed in the large intestine easily because of its lipophilicity, and then metabolized in the liver by O-methylation, glucuronidation, and/or sulfation (Bentz, 2017). We assessed if quercetin could be an active anti-sickling agent as suggested by Muhammad et al., (2019). However, we failed to demonstrate any appreciable anti-sickling activities for quercetin, thus, substitution of an alpha-L-rhamnosyl moiety at position 3 via a glycosidic linkage in quercitrin would be important for its biological activity. Confirmation of this mode of action could focus on use of X-ray crystallography of Hb-quercitrin to demonstrate this.

### Insights into the mechanism of quercitrin effects using metabolomics studies

To provide a wider appreciation of the effects of quercitrin we applied an -omic approach based on metabolite detection using direct infusion-high resolution mass spectroscopy. Multivariate statistical assessments of the derived data made a series of important observations. Firstly, it was apparent that even under normoxia the metabolomes of HbSS erythrocytes were unlike HbAA cells, and indeed, were more similar to anoxic cells. This could suggest that even under normoxia, HbSS cells are poised to sickle and indeed, the SEM of suggested cellular irregularities. However, the addition of quercitrin caused a shift in the HbSS metabolome so that it became similar to HbAA. This allowed us to assess the metabolomic difference between sickled and non-sickled erythrocytes that quercitrin was effectively correcting.

Detection of porphyrin metabolites was predictable as the breakdown of haemoglobin that is a feature of sickling (Odièvre et al., 2011). More mechanistic metabolite changes are suggested from the inositol phosphate metabolite, D-myo-inositol 1, 4, 5-trisphosphate (Ins(1,4,5) 3PO_4_). This inhibits human erythrocytes Ca^2+^-stimulatable, Mg^2+^-dependent adenosine triphosphatase (Ca^2+^-ATPase) activity (Davis et al., 1995) and the binding of calmodulin to the erythrocytes membrane (Berrocal et al., 2017). Such effects could perturb the Ca^2+^ activated Gardos channel leading to dehydration of HbSS-erythrocytes (Odievre et al., 2011). This suggestive data could be hinting at an additional role for quercitrin in influencing Ca^2+^ activated Gardos channels.

Another altered pathway involves bioenergetic metabolism. Glycerolipid pathways include monoacylglycerols (MAGs), diacylglycerols (DAGs), triacylglycerols (TAGs), phosphatidic acids (PAs), and lysophosphatidic acids (LPAs) with functions in energy generation (Zhang & Reue, 2017). The hexose monophosphate shunt which is parallel to glycolytic pathway and generates NADPH and pentoses as well as ribose 5-phosphate, a nucleotides-synthesis precursor (Jin & Zhou, 2019). Increased D-glucose 1-phosphate, when interconverted to D-glucose 6-phosphate, will also feed through to the glycolytic pathway involved in the generation of ATP in the anabolic generation of intracellular energy (Christodoulou et al., 2019). Such could indicate that the sickle cell could be exhibiting a tendency towards anaerobic respiration. Reduced glucose 1-phosphate levels in quercitrin-treated samples suggest that the anaerobic shifts are countered by quercitrin.

Sickle cells generate approximately two times more reactive oxygen species compared with normal red blood cells (Chirico & Pialoux, 2012) and this is linked to endothelial dysfunction, inflammation and multiple organ damage (Hundekar *et al*., 2010; Rusanova *et al*., 2010).

Decreased intravascular sickling has been linked with reduced oxidative stress and also an increased nitric oxide bioavailability (Dasgupta *et al*., 2010; Magalhães, 2011). Such oxidative effects with sickling were suggested from the metabolomic analysis as “ascorbate and aldarate” metabolism pathways were being affected in our experiments. It should be noted that glutathione, detected in our metabolome, also plays an important role in the anti-oxidative process (Kawasaki et al., 2005). Glutathione protects the red cells from oxidative damage, denaturation of haemoglobin (van ‘t Erve et al., 2013) and the formation of Heinz bodies, reduced cell deformability and intravascular haemolysis (Frewin, 2014).

## Conclusions

In this work, quercitrin was isolated from *A. cordifolia* and its anti-sickling activity was characterized. Quercitrin inhibits and marginally reverses HbSS erythrocytes under hypoxic conditions *in vitro*. Experimental evidence was presented to support its action against two features of SCA. Most importantly, quercitrin inhibited Hb polymerisation but it also stabilized HbSS-erythrocytes membrane to reduce erythrocytes fragility. Metabolomics was used to provide a wider description of the sickling process and the effects of quercitrin. All of these were reversed with quercitrin treatment.

## Author contribution statement

Conceptualization, O.A., A.W. and L.M.; methodology, OA., A.C., R.B., and S.B..; writing—original draft preparation, O.A.; writing—review and editing, all authors.; supervision, L.M. and A.W. ; project administration, L.M.; funding acquisition, L.M.. All authors have read and agreed to the published version of the manuscript.

## Acknowledgements

We appreciate the technical support provided by Dr. Manfred Beckmann and Ms. Helen Phillips with metabolomic profiling (Aberystwyth University UK).

