## Supplementary figures and tables for "Isolation and characterisation of quercitrin as a potent anti-sickle cell anaemia agent from *Alchornea cordifolia*"

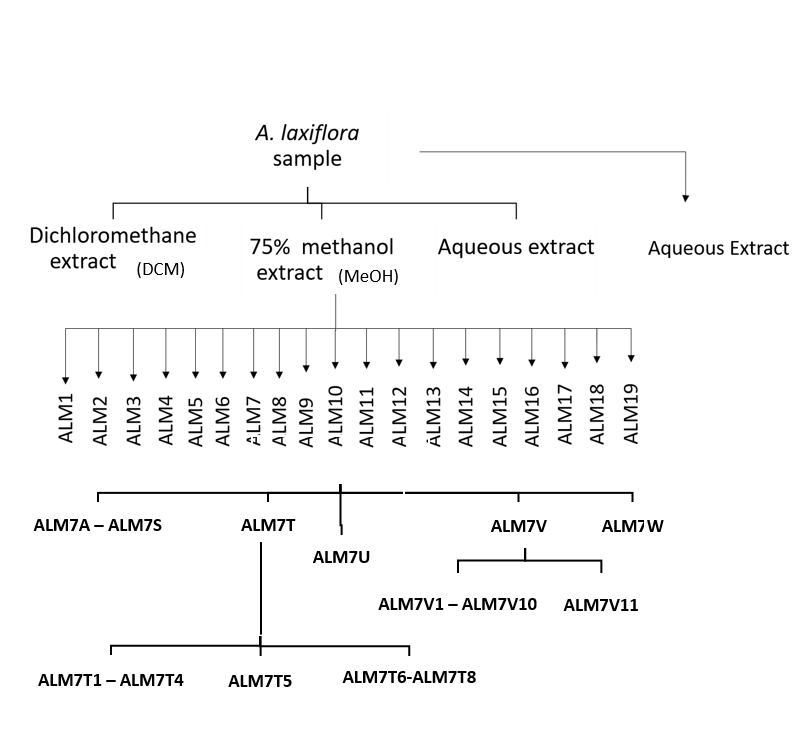

ALM7A – ALM7S

ALM7T

ALM7U

ALM7V

ALM7W

ALM7T1-ALM7T4

ALM7T5

ALM7T6-ALM7T8

Aqueous extract2

Aqueous extract1

**Fig. S1.** Schematic diagram for the purification of anti-sickling activities in *Alchornea spp.*

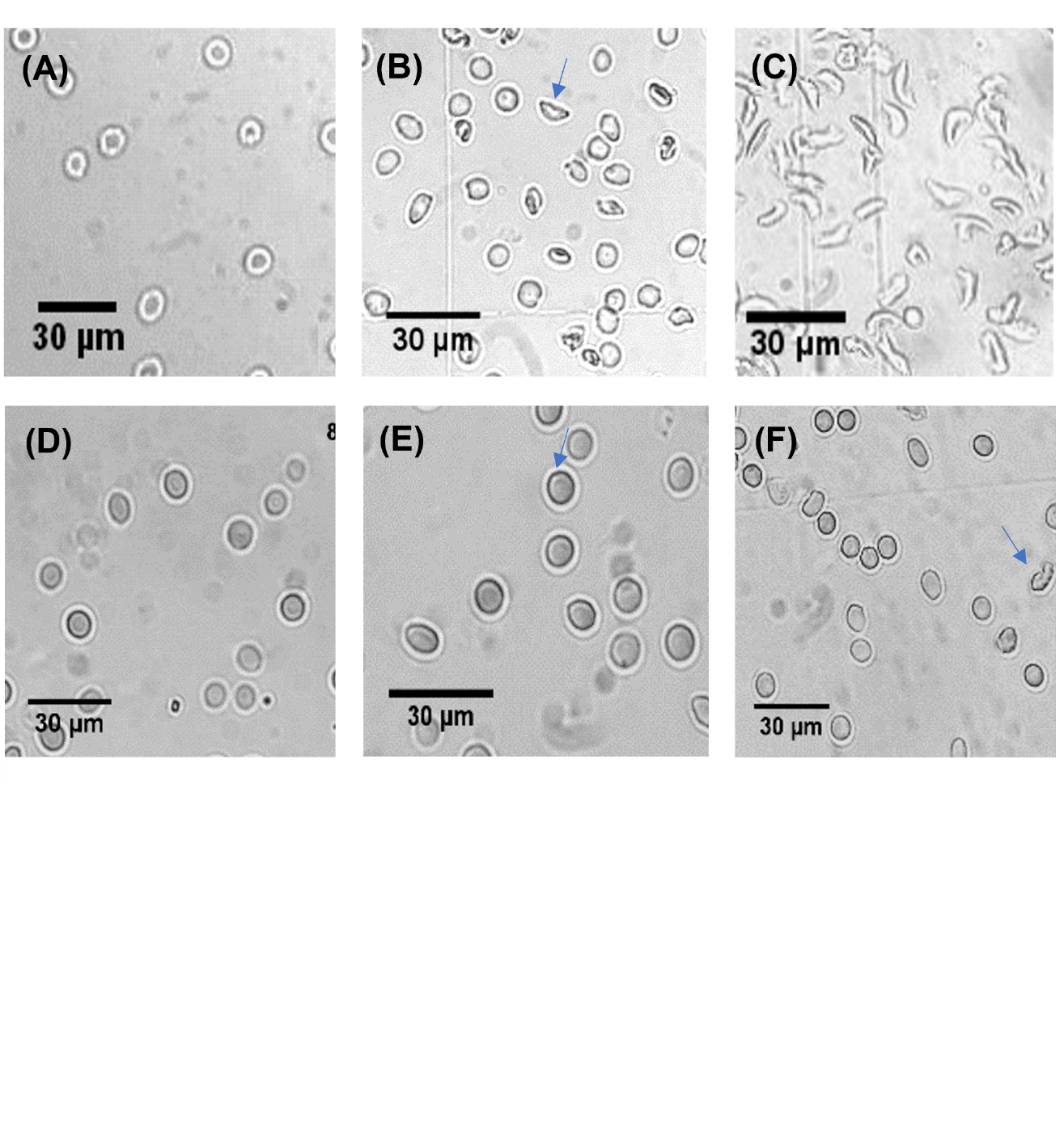

**Fig. S2** The rffect of *Alchornea spp* extracts and fractions on erythrocyte-sickling in Na_2_S_2_O_5_- induced hypoxia**.**

**(A)** HbAA normoxic **(B)** HbSS normoxic (sickled cell arrowed) **(C)** HbS hypoxic [hyp] (all cells sickled). **(D)** HbSS-hyp+ALM7T5 (0.4 mg/mL)**(E)** HbSS-hyp+ALM7T5 (0.2 mg/mL) **(F)** HbSS-hyp+ALM7T5 (0.05 mg/mL) (abnormal cells arrowed).

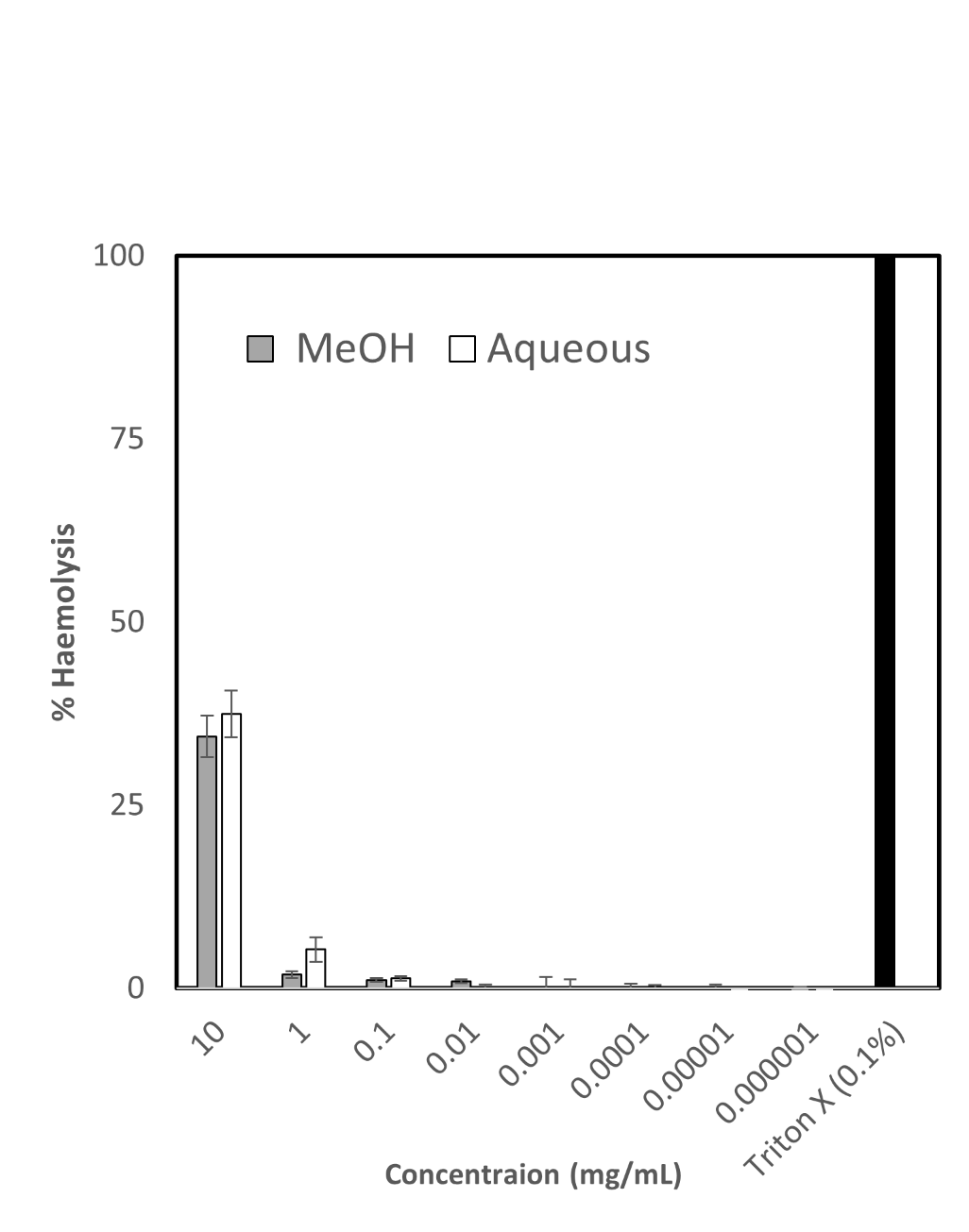

**Fig. S3.** Percentage haemolysis in HbSS RBCs incubated with various concentrations of MeOH and Aqueous extracts of *Alchornea spp* leaves compared with 0.1% Triton X (positive control with 100% haemolysis). *Typical data expressed as mean of experiments performed in triplicates

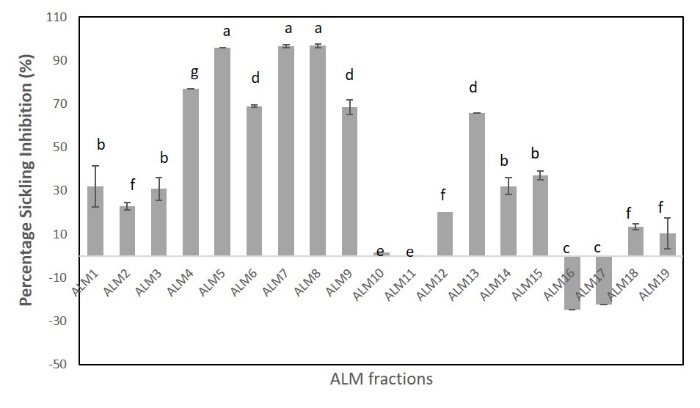

**Fig. S4.** The effect of ALM fractions (1 mg /mL) on erythrocytes-sickling under Na_2_S_2_O_5_- induced hypoxic condition.

Groups within which there is no difference are indicated by different letters. The data represents average of 3 similar results from repeat experiments.

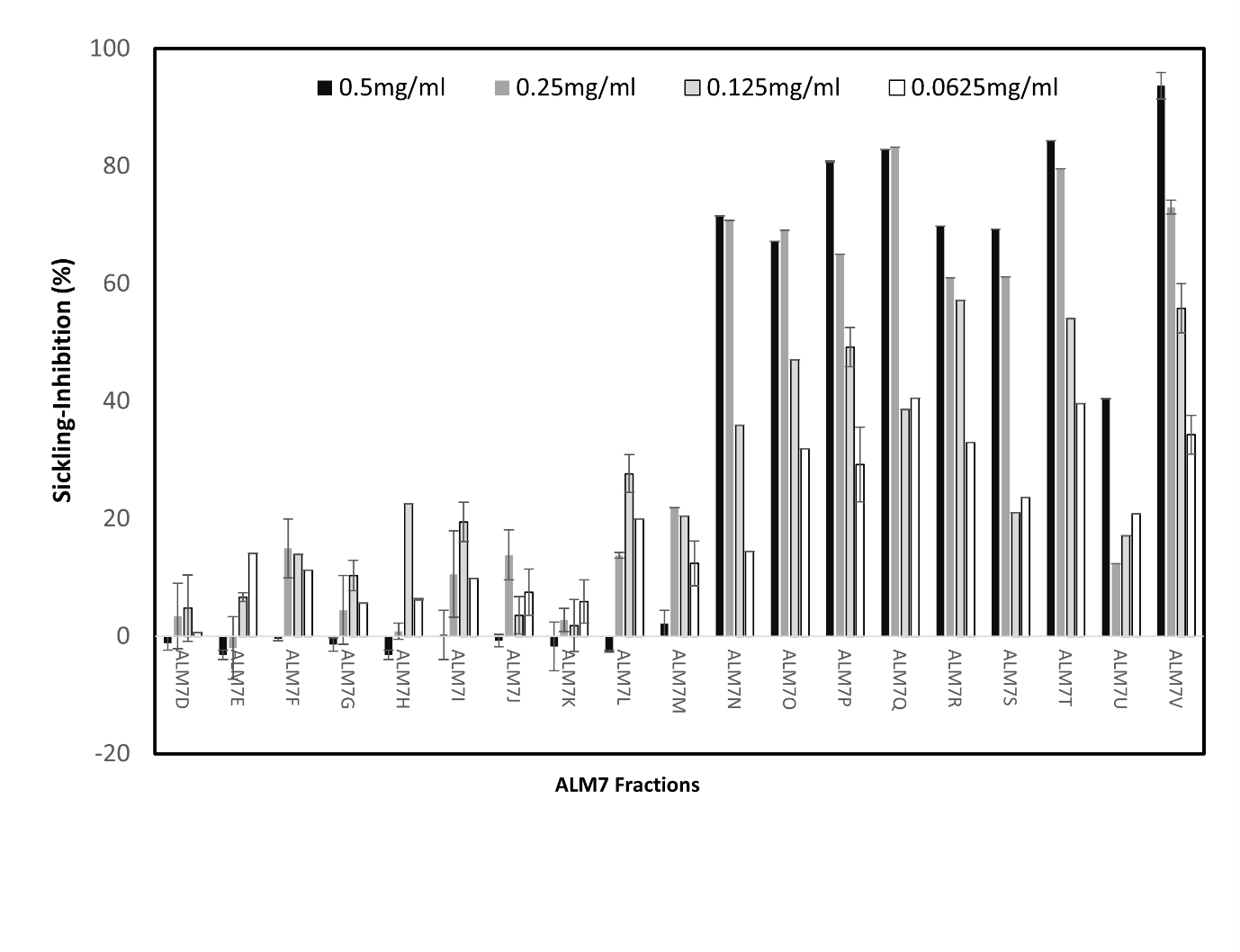
**Fig. S5.** Effect of ALM7 fractions (500 – 62.5 µg/mL) on erythrocytes-sickling under Na_2_S_2_O_5_- induced hypoxic condition.

Data represents average of 3 similar results from repeat experiments.

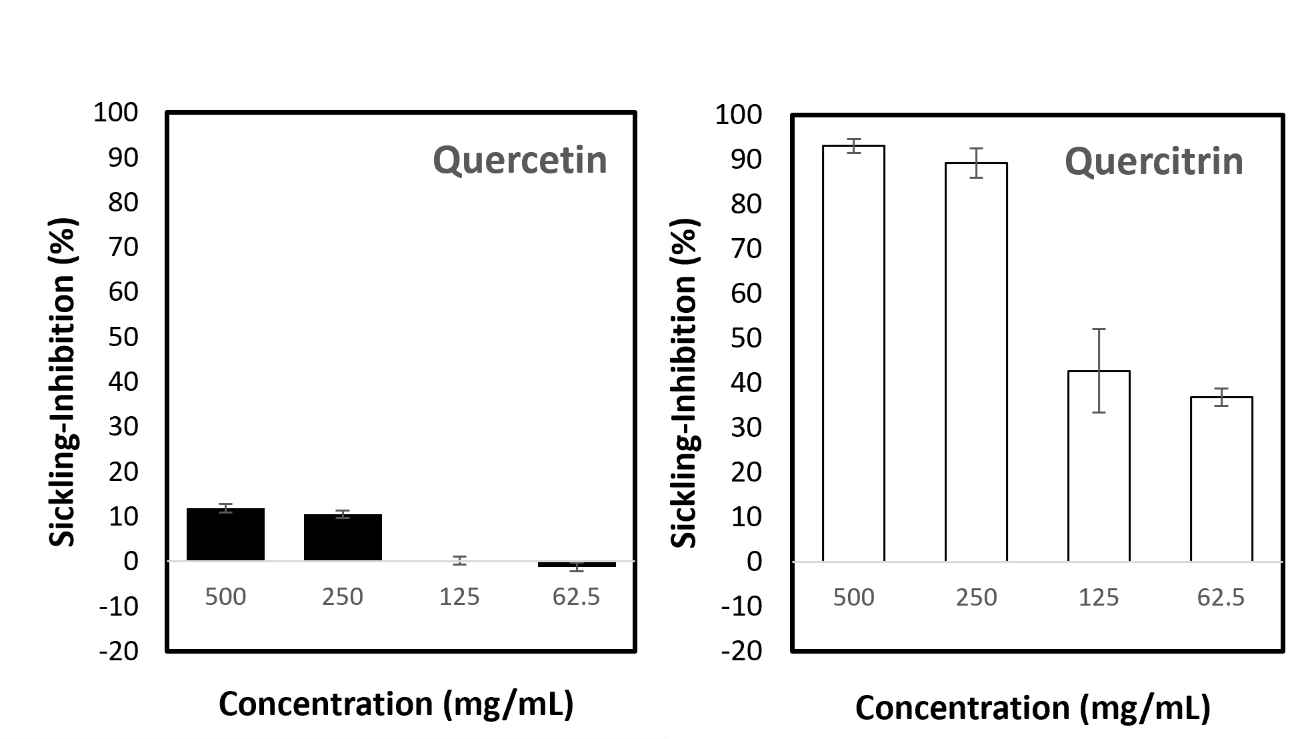

**Fig. S6.** Inhibitory Effects of Quercetin on HbSS erythrocyte sickling

*in vitro* assessment of erythrocyte sickling at low oxygen tension induced by Na_2_S_2_O_5_ after a 3 hour incubation period. These represents data obtained from three typical independent experiments.

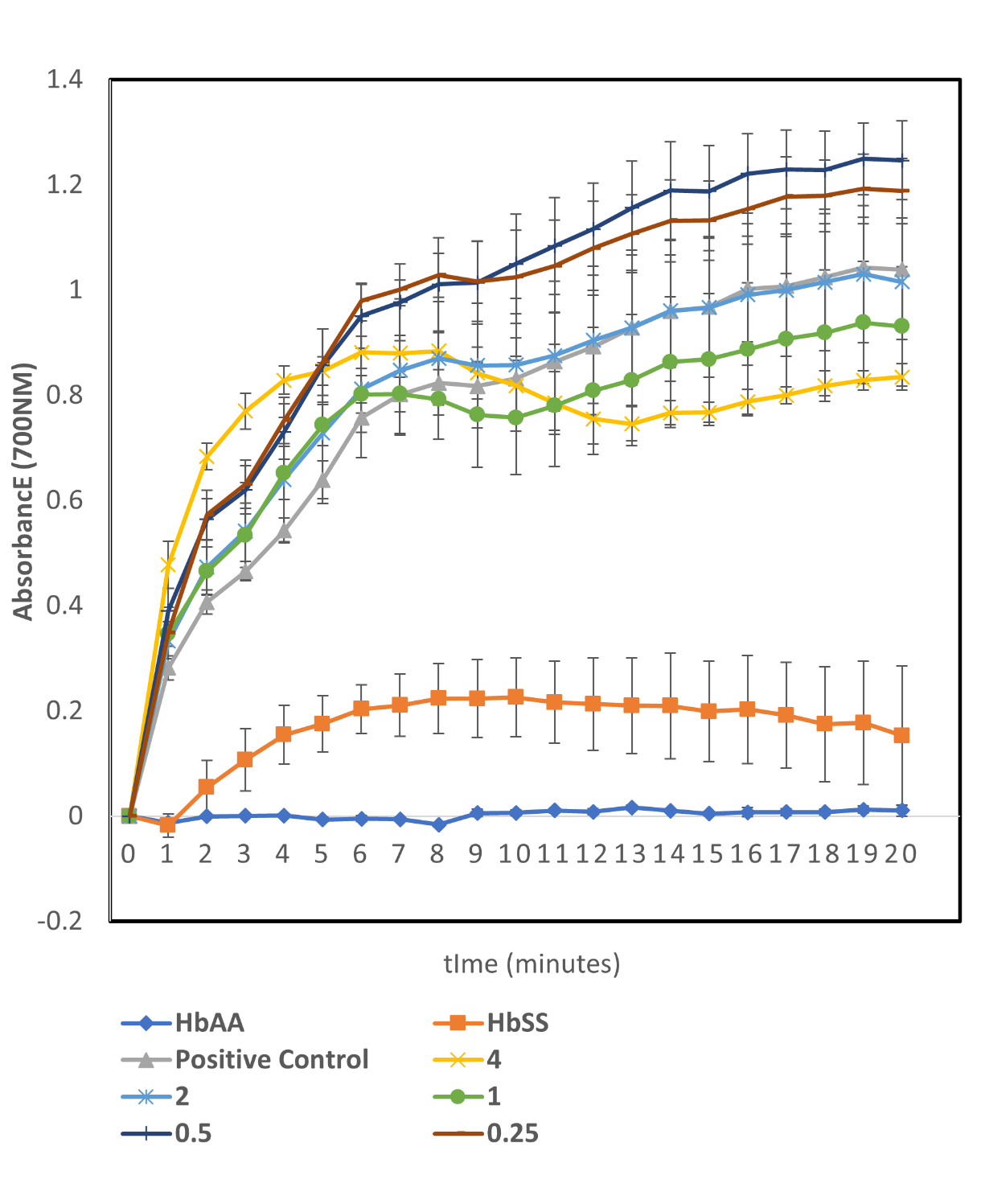

**Fig. S7.** Assessing the potential inhibitory Effects of quercetin on HbS Polymerisation.

Quercetin showed no inhibition of HbS polymerisation with absorbance of polymerised HbS similar to the positive control; deoxyHbS without treatment, at all assayed concentrations. DeoxyHbA (HbAA) was not polymerised as was effectively the negative control. OxyHbS (HbSS) was HbS not subjected to deoxygenation. This represents data obtained from three typical independent experiments performed in quadruples.

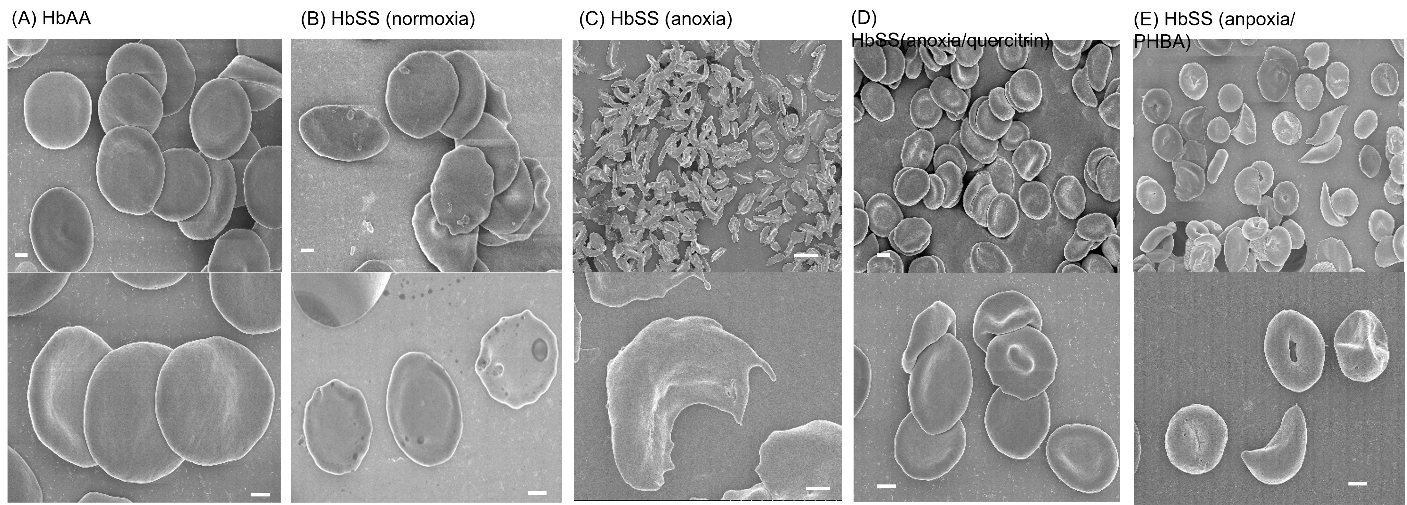

**Fig. S8.** The effect of *Alchornea* spp extracts and fractions on erythrocyte-sickling in N_2_- induced hypoxia

| **Pathway Names** | **Hits** | ***P* -value** |
| --- | --- | --- |
| Ascorbate and aldarate metabolism | 4 | 0.02452 |
| Cysteine and methionine metabolism | 8 | 0.0351 |
| Biosynthesis of unsaturated fatty acids | 8 | 0.0351 |
| Pentose and glucuronate interconversions | 7 | 0.03732 |
| Fatty acid biosynthesis | 2 | 0.04529 |
| Porphyrin metabolism | 2 | 0.04529 |

**Table S1.** : Pathway Analysis of *m/z* features significantly (P<0.05) different between the treated and untreated groups.
